# Genomics insights reveal multi-year maintenance of a new *Deltacoronavirus* infecting Seabirds from the Cagarras Island Archipelago Natural Monument, Brazil

**DOI:** 10.1101/2025.06.19.660416

**Authors:** Yago José Mariz Dias, Alexandre Freitas da Silva, Allan Rodrigues, Fernanda Gomes, Leonardo Corrêa, Paola Cristina Resende, Marilda Siqueira, Marina Galvão Bueno, Tatiana Ribeiro, Paula Gabriela Quintana, Larissa Schmauder Teixeira da Cunha, Luciana Reis Appolinario, Hudson Marques Paula Costa, Marcione Brito de Oliveira, Dilmara Reischak, Jose Reck, Gabriel Luz Wallau, Maria Ogrzewalska

## Abstract

Previous studies have identified various pathogens in seabirds, notably coronaviruses (CoVs) and influenza A viruses (IAVs), due to their potential to cause significant morbidity and mortality. The Cagarras Island Archipelago Natural Monument, located near Rio de Janeiro, Brazil, serves as nesting site for two species, the magnificent frigatebird (*Fregata magnificens*) and the brown booby (*Sula leucogaster*). Despite its ecological importance, no prior studies have investigated viral infections in these species, which share habitat interfaces with densely populated human areas. To address this gap, we sampled and tested seabirds for CoVs and IAVs from January 2022 to April 2024. Birds were captured and identified by species, age, and sex. Oropharyngeal and cloacal swabs, as well as blood samples, were collected. Viral RNA was extracted using the QIAamp Viral RNA Mini Kit, and the presence of IAVs was screened via real-time RT-PCR, while CoVs were screened using semi-nested RT-PCR. Sanger and metatranscriptomic sequencing were performed to identify viral strains and assess phylogenetic relationships. Of the 153 seabirds sampled, CoVs were detected in 6 individuals (9.1%) of *F. magnificens* and 16 individuals (18.4%) of *S. leucogaster*. No IAVs were found in either oropharyngeal or cloacal swabs, and all serum samples were negative for the presence of antibodies against the virus. We recovered two full deltacoronavirus genomes and eight additional draft genomes from *S. leucogaster* samples obtained from distinct sampling expeditions and additional enteroviruses, passeriviruses, and picornaviruses. Phylogenetic analysis revealed that the detected CoVs are closely related to avian deltacoronaviruses from environmental samples of *S. leucogaster* in the São Pedro and São Paulo Archipelago, indicating potential viral exchange between these seabird populations living at these distant islands. Moreover, multiple detections in different individuals at different time points are associated with specific Spike NTD deletions that have been shown to accumulate in immune escape lineages, supporting the long-term maintenance through new infections and reinfection of this virus in these bird populations. This is the first detection of CoVs in *F. magnificens*, highlighting their circulation in marine ecosystems. Further research is needed to understand the ecological and epidemiological implications, including potential cross-species transmission.

## INTRODUCTION

Seabirds comprise a vast diversity of species and include members of at least six avian orders, such as Sphenisciformes, Procellariiformes, Pelecaniformes, Suliformes, Phaethontiformes, and Charadriiformes, that share the characteristic of feeding at sea (Furness 1987). Most seabirds exhibit colonial breeding behavior, gathering in large numbers for extended periods each year to reproduce. Although seabirds show remarkable colony fidelity during the breeding season, they are also known to undertake extensive journeys during non-breeding periods to forage and explore potential future breeding sites (Furness 1987). This behavior highlights their potential role as important vectors in the dispersal of infectious agents across diverse ecosystems (McCoy, et al. 1999; Reed, et al. 2003; Altizer, et al. 2011; McCoy, et al. 2016). Among the viruses associated with seabirds, coronaviruses (CoVs) and influenza A viruses (IAVs) warrant special attention due to their potential to cause morbidity and, in some cases, mass mortality among both wild and domestic birds (Munster, et al. 2007; Muradrasoli, et al. 2010; Dusek, et al. 2014; Huang, et al. 2014; Lang, et al. 2016; Prosser, et al. 2022).

The Cagarras Islands Archipelago Natural Monument (MONA Cagarras) is an integral protection conservation unit established in 2010, with the primary purpose of preserving the ecosystem and serving as a nesting and refuge area for seabirds. Located in the municipality of Rio de Janeiro, approximately five km from Ipanema Beach, it harbors a sanctuary for seabirds, hosting one of the two main breeding colonies of magnificent frigatebirds (*Fregata magnificens*) in the South Atlantic and one of the largest colonies of brown boobies (*Sula leucogaster*) on the Brazilian coast (Cunha 2013) (**Suppl. Figure 1**). Along the Brazilian coast, smaller colonies of these two species can still be found, and they frequently follow fishing boats. In addition to their habit of catching prey near the water’s surface, frigatebirds often collect and bring marine debris, including trash, to their nests. Unfortunately, some individuals die after becoming entangled in fishing lines and other discarded gear (Cunha L. personal communication).

No previous study has been conducted in this area to investigate the presence of viruses. To address this research gap, we conducted a study focusing on collecting samples from seabirds to test for the presence of CoVs and IAVs, followed by metatranscriptomic sequencing of positive samples. By analyzing these samples, we aimed to gain valuable insights into the potential role of seabirds as hosts of these viruses within this unique environment - a massive colony of over 6,000 frigatebirds and approximately 2,500 brown bobbies.

## MATERIALS AND METHODS

### Study area and sample collection

The MONA Cagarras is composed of four islands (Cagarra, Redonda, Palmas, and Comprida) and two islets (Filhote da Cagarra and Filhote da Redonda). Two of them, the Cagarra and the Redonda Islands, serve as nesting sites for brown bobbies and magnificent frigatebirds, respectively (**Figure 1**). The Cagarra Island (23° 1’36.86”S, 43°11’33.17”W) covers an area of approximately 9.3 hectares. The island’s vegetation is low-lying and shrubby, with patches of bare soil that harbor important nesting grounds for brown boobies. The Redonda Island (23° 4’14.39”S, 43°11’41.43”W), on the other hand, is the largest and tallest among the islands of the archipelago with 41,4 hectares of emerged area, located 8 km from the coast. It features a rocky formation with cliffs from the base to the summit. The island vegetation is composed of herbs and small shrubs in the lower regions and a compact, low forest at its summit. The Redonda Island stands out in the sea and hosts a nesting colony of magnificent frigatebirds, where approximately 2,640 active nests are estimated. Brown bobbies reproduce on the Redonda as well, but in lower densities (Bovini, et al. 2013; Cunha 2013; Moraes and Seoane 2013).

**Figure 1.**
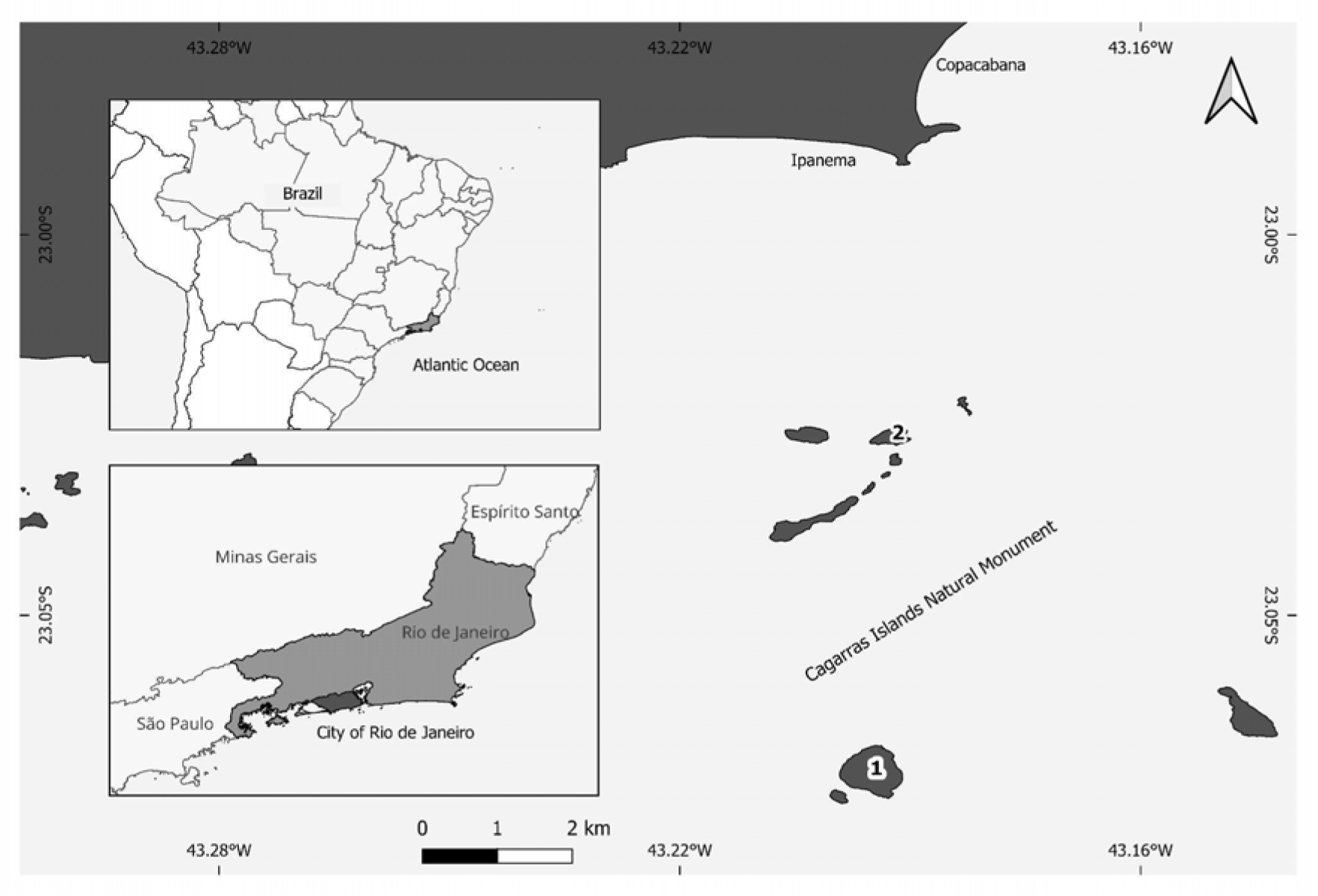
Localization of the MONA Cagarras and the island where the birds were sampled: 1 – Redonda Island, 2 - Cagarra Island (Source: Generated by authors using QGIS 3.32.2 based on public governmental shapefiles).

The MONA Cagarras also protects other bird’ species such as kelp gulls (*Larus dominicanus*), which reproduce on the islands, and other species such as black vultures (*Coragyps atratus*), American oystercatcher (*Haematopus palliatus*), and black-crowned night-heron (*Nycticorax nycticorax*), which are frequently visiting the islands (Cunha 2013).

### Biological sample collection

Field collection expeditions were conducted from February 2022 to April 2024 at monthly intervals. Birds were selected by convenience, aiming to include a variety of ages and an equal number of each species. All birds were captured using a net (3 cm mesh size, 40 cm diameter). After being identified to the species level, they were classified according to sex and age class (chick, immature, and adult) (ICMBio 2018). Adult birds were banded on their legs with metal bands provided by the National Center for Research and Conservation of Wild Birds (CEMAVE). Following identification, oropharyngeal and cloacal samples were collected using sterile Rayon swabs, which were then placed into cryotubes containing 1 mL of Viral Transport Medium (VTM). Additionally, 2-3 mL of blood was collected from the medial metatarsal vein using a 3-mL syringe and a 13 × 4.5 mm needle. All samples were kept refrigerated for up to five hours and then brought to the laboratory, where swabs were frozen at -80°C for later analysis. Blood samples were centrifuged to obtain sera, which were then frozen at -20°C.

### Viral RNA extraction

In the laboratory, viral RNA extractions from individual oropharyngeal and cloacal swab samples were performed using the QIAamp Viral RNA Mini Kit (Qiagen, CA, USA). A final elution buffer volume of 60 µL was obtained as recommended by the manufacturer. The isolated RNA was immediately stored at -80°C for molecular analysis. For each extraction step, RNase/DNase-free water was used as a negative control.

### Screening for Influenza A Viruses (IAVs) by Real-Time RT-PCR

Molecular detection of IAVs was performed using real-time PCR (rRT-PCR), employing a set of oligonucleotides InfA-F 5’-GAC CRA TCC TGT CAC CTC TGAC-3’, Infa-R 5’-AGG GCA TTY TGG ACA AAK CGT CTA-3’, and probe Infa-P 5’-TGC AGT CCT CGC TCA CTG GGC ACG-3’, labeled with FAM at the 5’ end and BHQ-1 at the 3’ ends. The reaction targeted the matrix gene, following the protocol established by the Collaborative Influenza Center, Centers for Disease Control and Prevention, Atlanta, GA (Shu, et al. 2011) for all subtypes of influenza A. Results were analyzed using SDS Software v1.4, as described by (Shu, et al. 2011). Samples with a Cycle threshold (Ct) value below 40 and exhibiting a characteristic sigmoidal curve were considered positive.

### Coronavirus (CoVs) screening by conventional pancoronavirus RT-PCR

Coronavirus detection was performed using conventional semi-nested RT-PCR for pan-coronavirus detection, following the protocol developed by (Chu, et al. 2011). RNA was subjected to RT-PCR targeting a 620-base pair fragment of the RNA-dependent RNA polymerase (RdRp) gene. In brief, RdRpS1 (5’-GGKTGGGAYTAYCCKAARTG-3’) and RdRpR1 (5’-TGYTGTSWRCARAAYTCRTG-3’) primers were used in the first reaction with the One-Step RT-PCR Enzyme Mix (Qiagen) kit, followed by Bat1F (5’-GGTTGGGACTATCCTAAGTGTGA-3’) and Bat1R (5’-CCATCATCAGATAGAATCATCAT-3’) primers in the second PCR with the Phusion High-Fidelity PCR kit (Thermo Fisher), amplifying a 440 base pair product. Cycling conditions followed recommendations of the OneStep RT-PCR kit (Qiagen). They were conducted on a Veriti Thermal Cycler (Applied Biosystems), including reverse transcription (50°C, 30 min), reverse transcriptase inactivation and DNA polymerase activation (95°C, 15 min), followed by 40 cycles of DNA denaturation (94°C, 45 s), annealing (52°C, 45 s), and extension (72°C, 45 s), with a final extension step (72°C, 10 min). Subsequently, conditions for the second PCR using the Phusion RT-PCR Enzyme Mix (Sigma-Aldrich) were as follows: initial denaturation (98°C, 30 s), followed by 35 cycles of DNA denaturation (98°C, 15 s), annealing (52°C, 15 s), extension (72°C, 30 s), and a final extension step (72°C, 5 min). The amplified products (440 bp) were visualized on a 1.5% agarose gel and purified using the QIAquick Gel Extraction Kit (Qiagen) following the manufacturer’s recommendation.

### Sanger sequencing

The Sanger sequencing reaction was prepared using BigDye Terminator v3.1 Cycle Sequencing Kit (Life Technologies) with primers Bat1F and Bat1R. The sequencing was performed using the ABI 3730 DNA Analyzer (Applied Biosystems).

### Metatranscriptomic sequencing

After validation of deltacoronavirus detection in samples by Sanger sequencing, we processed the same samples by metatranscriptomics to recover the full deltacoronavirus genomes. Positive samples were treated with the Ambion® TURBO DNA-free™ Kit (Invitrogen) to remove residual genomic DNA from the extracted RNA. Initially, 30 µL of RNA was combined with 3 µL of 10× TURBO DNase Buffer and 1 µL of TURBO DNase in a microcentrifuge tube. The mixture was incubated at 37°C for 30 minutes to allow DNase to degrade any contaminating DNA. Following incubation, 3 µL of Stop Solution was added, and the sample was further incubated for 5 minutes at room temperature to ensure complete inactivation of the DNase. Samples were then centrifuged at 10,000 × g for 1.5 minutes, and the supernatant was transferred to a new tube. Host rRNA was depleted using the Illumina Ribo-Zero Plus rRNA Depletion Kit (Illumina, San Diego, CA, USA). The resulting RNA was used for library preparation with the Illumina Stranded Total RNA Prep, Ligation with Ribo-Zero Plus kit (Illumina, San Diego, CA, USA), which includes enzymatic fragmentation, adapter ligation, and strand-specific cDNA synthesis. Libraries were sequenced on the Illumina® MiSeq platform using a 300-cycle Micro cartridge for an initial test run and subsequently on the NextSeq platform (Illumina, San Diego, CA, USA) using a NextSeq 1000/2000 P2 cartridge. To enhance genome coverage, sequencing reads from both MiSeq and NextSeq platforms were combined for the final analysis.

### Metatranscriptome assembly and extension

Raw reads were first analyzed by quality using fastp v0.23.4 (Chen, et al. 2018) and trimmed using a cutoff of 30 for the Phred score. Trimmed reads were assembled by the *de novo* approach using MEGAHIT 1.2.9 (Li, et al. 2015). The assembled contigs were analyzed using DIAMOND (Buchfink, et al. 2015) running a blastx mode against the viral Refseq database recovered on May 5th, 2024. The sequences showing the five best hits on viral sequences were submitted to a reciprocal Diamond blastx analysis against the non-redundant (NR) database from NCBI to remove false negative hits. The contigs showing hits against viral proteins in these two analyses were kept and further analyzed. Viral contigs were assessed on the ViralComplete (Antipov, et al. 2020) tool to classify them into full-length or partial viral sequences and to recover the proteins that were predicted by Prodigal (Hyatt, et al. 2010). Partial sequences with the same taxonomic classification were retrieved from the following samples: CA32SR, CA43LC, CA42LC, CA87F, CA121SR, and CA124SR, and used for contig extension. Full-length contigs (CA_50SR_PBS_puro_combined_k141_2614, CA100SR_PBS_puro_combined_k141_1923 for *Deltacoronavirus* and CA100SR_PBS_puro_combined_k141_854 for *Enterovirus*) were then used as references in Multi-CSAR (Liu, et al. 2022) to reorder the fragmented contigs and insert “N” regions where necessary. The lengths of these “N” regions were verified and corrected by aligning the references, fragmented contigs, and Multi-CSAR outputs using MAFFT v7.511. The alignments were visualized and manually inspected in AliView (**Suppl. Figure 2)**. After extension, only contigs longer than 600 bp were retained for downstream analysis. Coverage metrics were obtained and calculated using the coverM tool (https://github.com/wwood/CoverM).

### Phylogenetic analysis

For the phylogenetic analysis, we used NCBI Virus to recover the closest sequences from NCBI in relation to the identified viruses. For this, we used the largest contig or RdRp amino acid sequences from each virus group identified as a query on BLAST analysis implemented through the NCBI virus platform. We conducted nucleotide or amino acid phylogenetic analysis depending on the identity of the viruses identified relative to sequences from the databases. The recovered sequences and those identified in this study were aligned using MAFFT v7.511, then visualized, edited, and inspected in Aliview (Larsson 2014). In certain cases, alignments were trimmed using the Trimal tool (Capella-Gutierrez, et al. 2009). The Maximum Likelihood phylogenetic analysis was performed using IQ-TREE 2.3.6 (Minh, et al. 2020) with the SH-aLRT test with 1000 replicates and ultrafast bootstrap with 10000 replicates applying the NNI (Nearest Neighbor Interchange) search. The best evolutionary models were selected by ModelFinder (Kalyaanamoorthy, et al. 2017). The trees were visualized and annotated using the ggtree R package (Yu, et al. 2017) and Figtree (http://tree.bio.ed.ac.uk/software/figtree/).

### Serology testing

Serum samples were tested for the presence of antibodies against Influenza A using a competitive ELISA kit (Inf A Multi ELISA CK401 BioCheck, BioCheck BV, Reeuwijk, The Netherlands).

### Statistical analysis

The Chi-square test of independence (McHugh 2013) was employed to evaluate the association between categorical variables and the presence of coronavirus. The test was applied to data across different variables, including species, age, sex, and sample type, to determine whether any significant relationships existed between these factors and the occurrence of the virus. For individual species (*S. leucogaster* and *F. magnificens*), the same test was conducted separately, focusing on the relationship between age, sex, sample type, and coronavirus presence. All analyses were performed on the R platform version 4.4.0 (Team 2024).

## RESULTS

### Species sampling

A total of 153 seabirds were captured and sampled at the MONA Cagarras between January 2022 and April 2024, including 66 *F. magnificens* (29 chicks, 16 immatures, 11 females, 10 males) and 87 *S. leucogaster* (34 chicks, 14 immatures, 24 females, 15 males).

### CoVs and IAVs screening

No IAVs were detected in 153 bird species from either oropharyngeal or cloacal swabs. Additionally, all 144 serum samples collected and tested were negative for the presence of antibodies against the virus.

On the other hand, Coronavirus RNA was detected in both *F. magnificens* and *S. leucogaster*, with positive individuals recorded in multiple sampling periods between 2022 and 2024. Among *F. magnificens*, six individuals (9.1%) tested positive, including both chicks and adults, suggesting potential intra-species circulation over the whole year. Similarly, 16 individuals of *S. leucogaster* (18.4%) were positive, also spanning different age groups and sampling dates (**Figure 2, Suppl. Tables 1 and 3**). We obtained high-quality partial RdRp sequences from twenty-one out of 26 positive samples (**Figure 2, Suppl. Table 1 and 3).**

**Figure 2.**
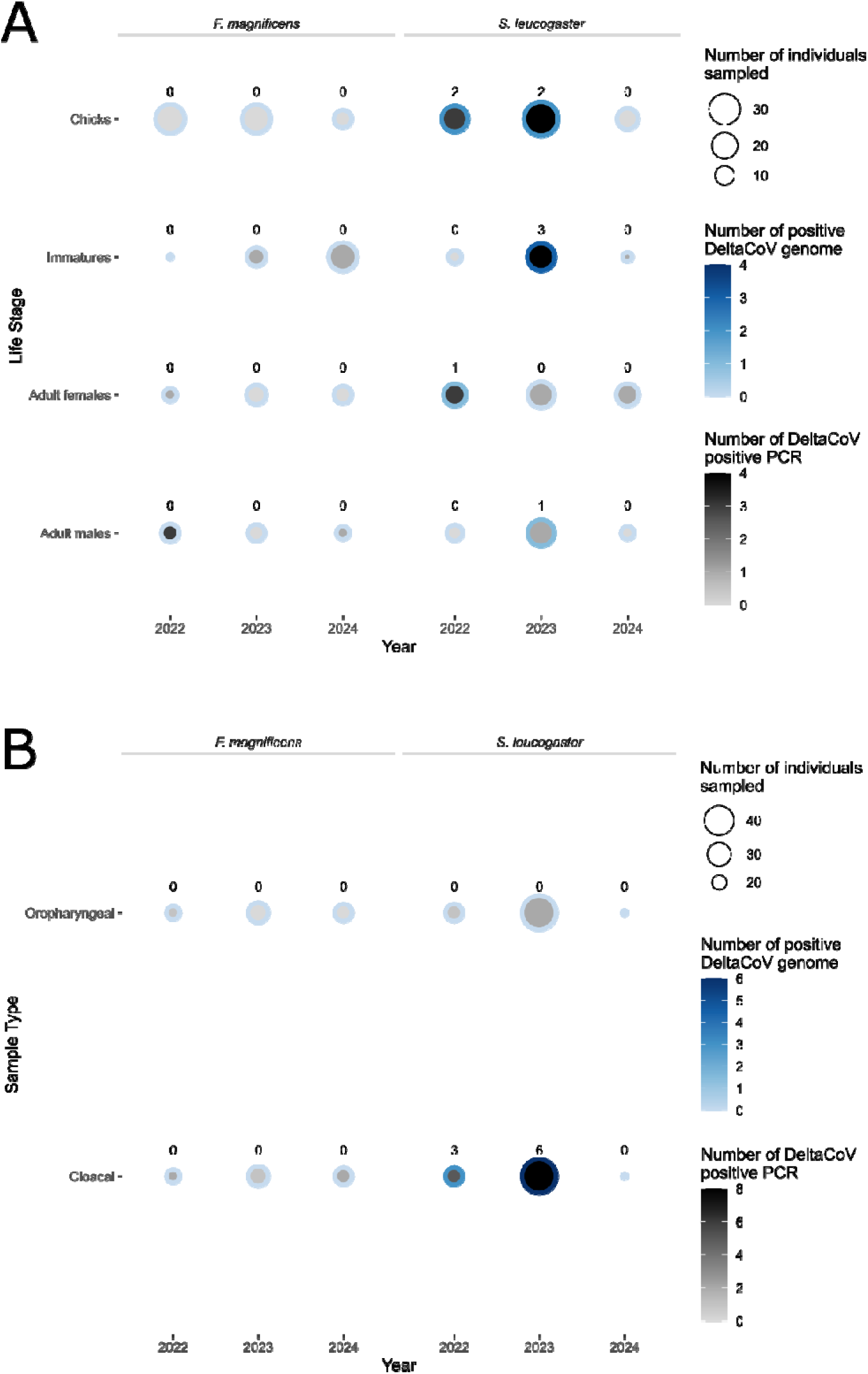
Detection by PCR and genome sequencing by life stage and sample type. The graph shows coronavirus detection counts by species by (A) life stage sex, and (B) sample type.

RdRp fragments obtained by Sanger sequencing were used as queries to recover homologous sequences from the databases, nucleotide sequence identities between the sequences recovered in this study and the sequences from the database ranged from 94,6% to 100%, showing a close relation with avian deltacoronaviruses previously identified in feces of *S. leucogaster* from the Sao Pedro Sao Paulo Archipelago (SPSP) (Gomes, et al. 2025). The DeltaCoV phylogenetic reconstruction confirmed the relationship of the deltacoronaviruses here characterized with those from SPSP forming a distinct clade (monophyletic group) with high ultrafast bootstrap and SH-aLRT support, indicating that the viruses isolated from these birds shared a recent common ancestor (**Suppl. Figure 3**).

The Chi-square test of independence was used to assess the relationship between variables (i.e, species, sex, lifestage, sample type) and the presence of the DeltaCoVs. No significant associations were found across all variables tested (**Suppl. Table 2**).

### Metatranscriptome analysis

To more in depth characterize the DeltaCoV found we performed metatranscriptomic sequencing of twenty-seven PCR-positive and Sanger validated samples from two species: *S. leucogaster* (CA28SR, CA29LC, CA32LC, CA34LC, CA34SO, CA34SR, CA42LC, CA43LC, CA50SR, CA80SR, CA87F, CA99LC, CA100SR, CA153F, CA153SR and CA155F) and *F. magnificens*: (CA33LC, CA33SR, CA35LC, CA35SO, CA40SO, CA51SR, CA51LC, CA121SR, CA124SR, CA159F and CA159SR). Total reads per sample ranged from 2.8 to 218 million raw reads (**Suppl. Table 4**). We identified a total of 40 viral contigs that ranged in size from 604 to 26,189 bp, representing fragmented or complete viral genomes (**Suppl. Table 4**). Our analysis revealed the presence of viral sequences belonging to five different viral families, such as *Picornaviridae, Picobirnaviridae, Nodaviridae, Coronaviridae,* and *Caliciviridae* (**Figure 3** and **Suppl. Table 4**). Different viral taxa were identified in samples from *S. leucogaster* (*Caliciviridae, Deltacoronaviruses, Passerivirus, Picobirnavirus, Enteroviruses, Hepatoviruses,* and others as unknown genus or family). In contrast, only *Picornaviridae,* such as enteroviruses and other contigs classified as unknown, were identified from *F. magnificens* samples (**Suppl. Table 4**).

**Figure 3.**
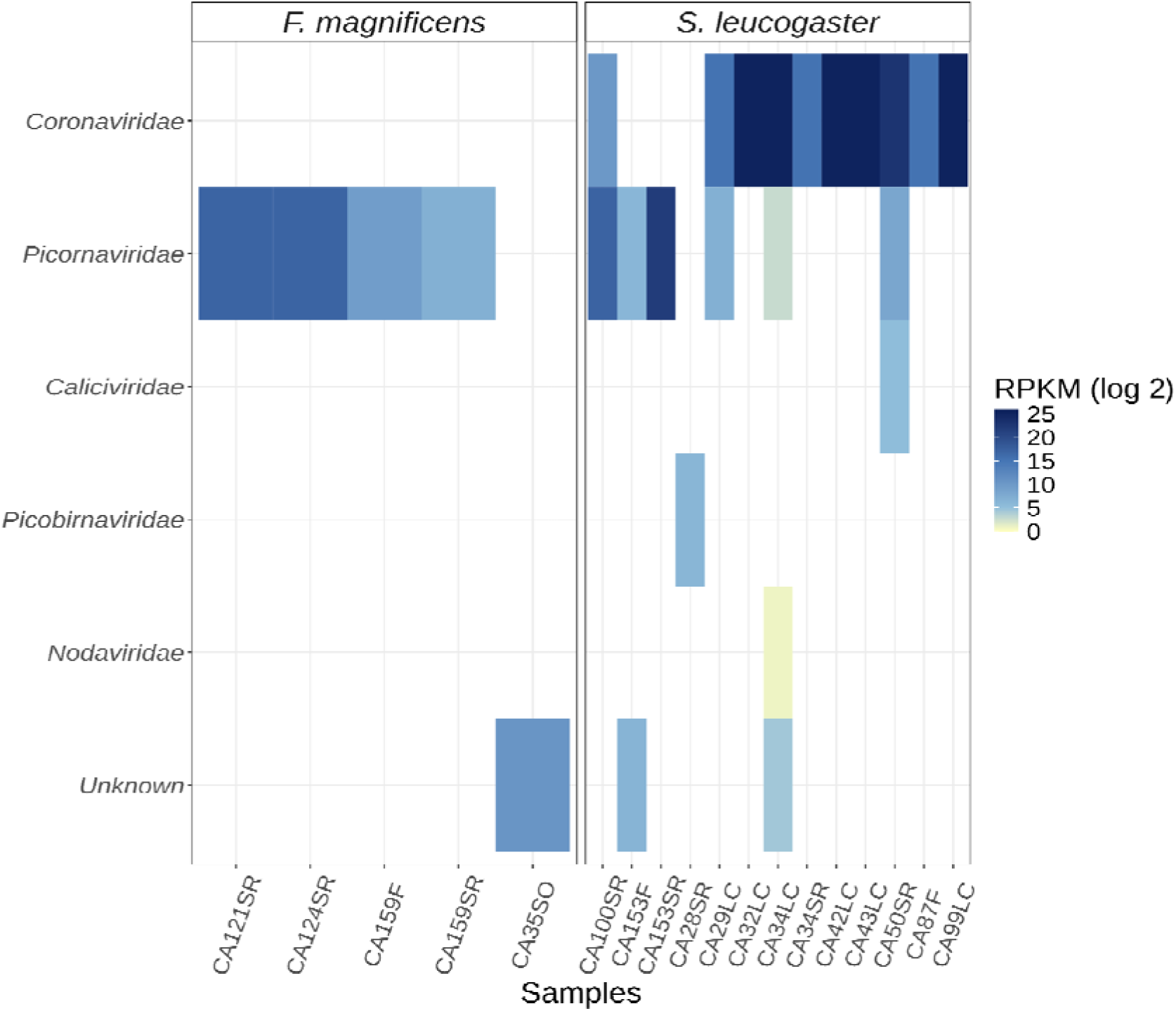
Heatmap displaying reads per kilobase per million mapped reads (RPKM) of *Picornaviridae, Picobirnaviridae, Nodaviridae, Coronaviridae, Caliciviridae,* and viral families with no current formal classification (Unknown) within each sample.

Ten *Coronaviridae* sequences were recovered from cloacal swabs, cloacal wash, and feces samples of *S. leucogaster* based on metatranscriptomic sequencing (CA50SR, CA100SR, CA29LC, CA32LC, CA34LC, CA34SR, CA43LC, CA99LC, CA87F, and CA42LC) (**Figure 3** and **Suppl. Table 4**). Of those, seven were complete or almost complete genomes (**Figure 4** and **Suppl. Table 4**). Deltacoronavirus genomes from *S. leucogaster* showed a high abundance (RPKM values ranging from 37.14 to 38,217,535.00), representing from 2.32% to 11.11% of passing filter reads (**Suppl. Table 4**). This deep sequencing coverage allowed us to recover complete genomes with a maximum coverage depth of 20,697.41x from the CA100SR sample (**Suppl. Table 4**). Our results revealed that these sequences were similar to Chinese pond heron coronavirus XN11 based on DIAMOND blastx analysis with best hit in ORF1a (WOR09770.1 and WOR09765.1), M protein (WOR09768.1) and N protein (WOR09769.1), showing an amino acid similarity of 63-68%, 88% and 53%, respectively (**Suppl. Table 4**). Regarding *F. magnificens* samples, we did not identify any coronavirus contigs.

**Figure 4.**
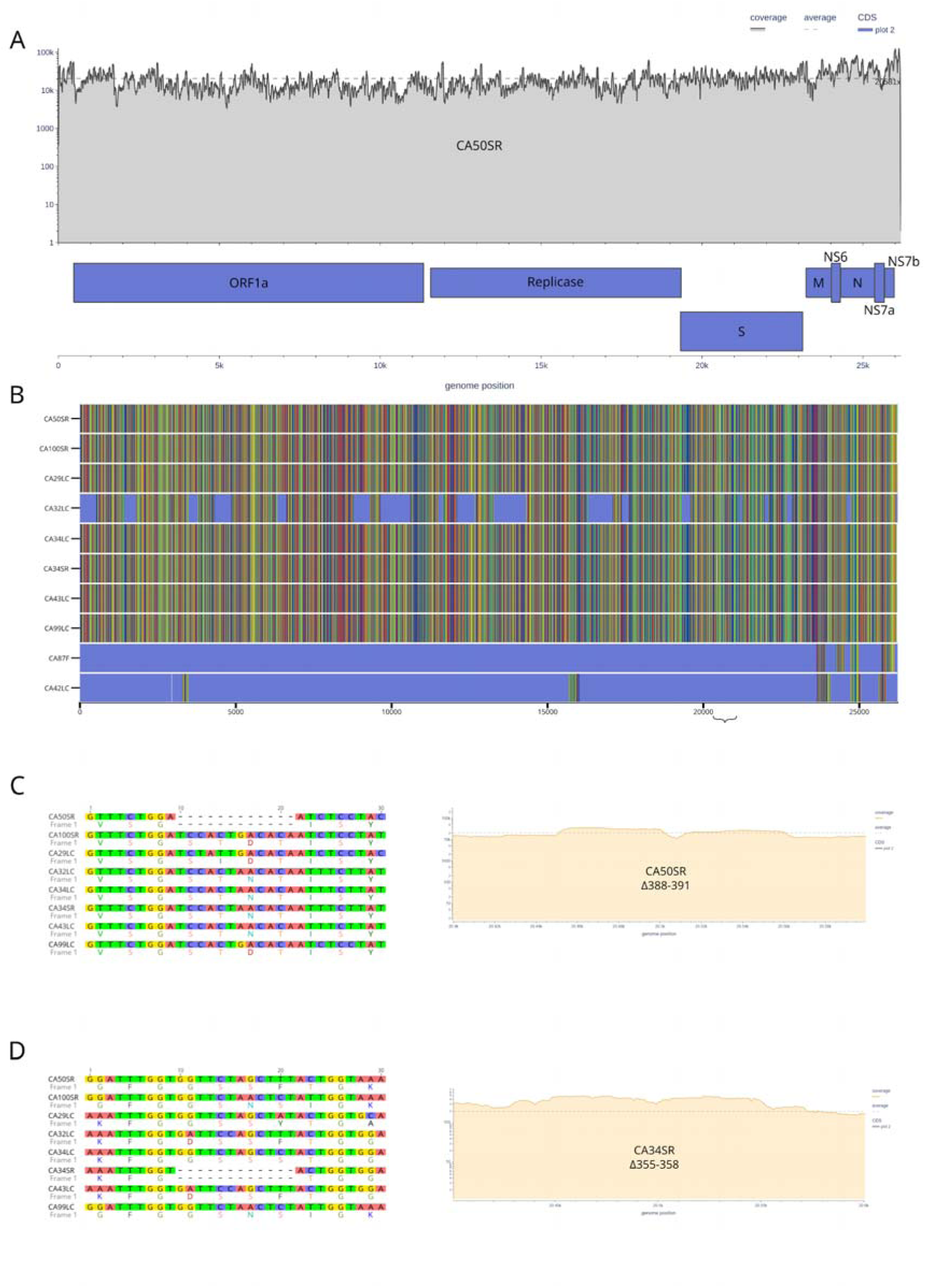
A) Coverage plot of the DeltaCoV genome from the CA50SR sample, generated using BAMdash. B) Multiple sequence alignment of all contigs identified as deltacoronavirus. Blue regions indicate stretches of “N”s resulting from the scaffolding process. C) Left: Multiple alignment highlighting a 12-nucleotide in-frame deletion in the spike coding region. Right: Coverage plot of the same contig. D) Left: Multiple alignment highlighting a 12-nucleotide in-frame deletion in the spike coding region. Right: Coverage plot of the same contig.

Two of the Delta-CoV genomes (CA50SR, Δ388-391, and CA34SR, Δ355-358) carry 12 nucleotides in frame deletions in different positions of the Spike protein that lead to 4 amino acid removal from the N-terminal region (**Figure 4**). Moreover, another in-frame deletion of 12 nucleotides and 4 amino acids was also detected in the ORF1a (RdRp N-terminal region) region (**Suppl. Figure 4**). These three deletions were well supported by reads (**Figure 4** and **Suppl. Figure 4**).

In addition, we identified another 30 viral contigs representing sequences with similarity in the *Durnavirales, Nodamuvirales,* and *Picornavirales* order (**Suppl. Table 4**). Thirteen contigs from *Picornaviridae* were assigned within three distinct genera such as *Enterovirus* genus (4), *Hepatovirus* (3) and *Passerivirus* (6) for samples from *S. leucogaster* (11) and *F. magnificens* (2) that showed similarities ranging from 26% to 96% with previously characterized enteroviruses, 25% to 43% with *Hepatovirus* and 33% to 68% with *Passerivirus*. Six contigs were assigned to the *Passerivirus* genus for *S. leucogaster* (CA50SR, CA100SR, and 29LC) and showed 33 to 68% similarity to known *Passerivirus* sequences. Two *Picobirnaviridae* sequences were identified and assigned to the *Picobirnavirus* genus.

Phylogenetic reconstruction of the *Deltacoronavirus* genus positioned the ten new genomes identified in *S. leucogaster* within the subgenus *Herdecovirus* (**Figure 5A**). These genomes clustered together with partial RdRp *Deltacoronavirus* sequences previously characterized from feces of the same species sampled in 2022 at the São Pedro and São Paulo Archipelago (**Figure 5B**). The enterovirus phylogenetic tree revealed that the enterovirus sequences identified in *S. leucogaster* and *F. magnificens* clustered within a clade with sequences from chimpanzees and humans (**Suppl. Figure 5**). One enterovirus from *Pan troglodytes troglodytes* was the basal branch, while the other enteroviruses were identified here. For sequences identified within the *Passerivirus* genus, we reconstructed one phylogenetic tree including the most similar sequences from the *Picornaviridae* family (**Suppl. Figure 6**). Sequences from *S. leucogaster* clustered within a clade that was positioned together with other passeriviruses from passerine birds such as thrushes *Uraeginthus granatina, Turdus pallidus,* and *T. hortulorum* (**Suppl. Figure 6B**). This passerivirus clade was positioned as a sister group of the *Sicinivirus* genus clade (**Suppl. Figure 6**). For *Caliciviridae* sequences, we reconstructed a phylogenetic tree including only one amino acid sequence representing a fragment of RdRp identified from *S. leucogaster* data. We further analyzed phylogenetically this sequence, and we named it as *Picorna*-like virus CA50 (**Suppl. Figure 7**). The *Picorna*-like virus CA50 was positioned as a basal branch of a clade containing other viruses such as Cryptolin_calicivirus_1 (WOC29232.1), Beihai_sesarmid_crab_virus_2 (YP_009333602.1), Wenzhou_picorna-like_virus_38 (APG78569.1), and Rudphi_virus_1 (AYN75566.1) (**Suppl. Figure 7**). These sequences were identified in other sea invertebrates such as clams *Ruditapes philippinarum, Gastropoda,* and crabs *Sesarmidae* (**Suppl. Figure 7**).

**Figure 5.**
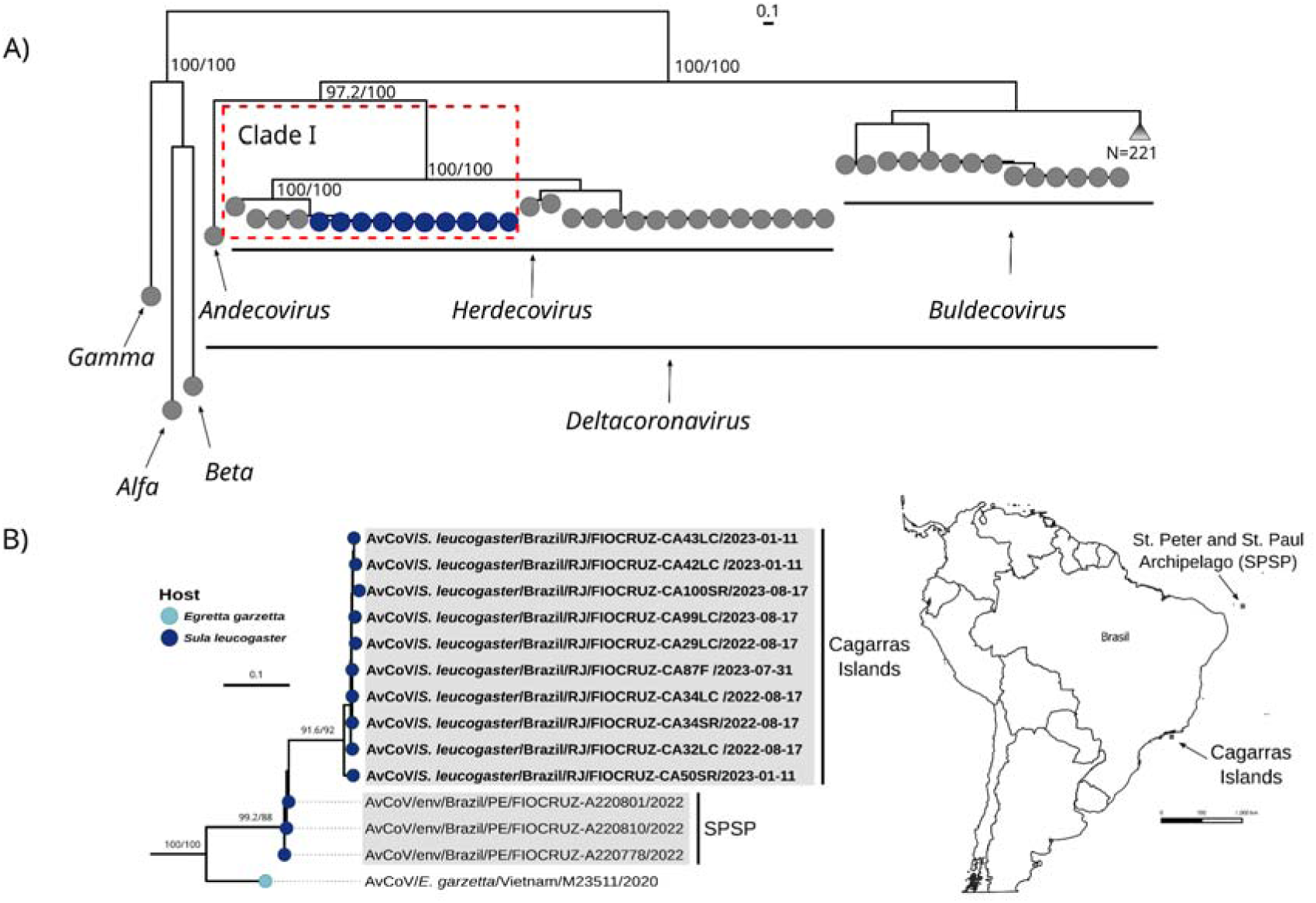
Phylogenetic analysis of deltacoronaviruses identified in *S. leucogaster.* A) Phylogenetic analysis of 337 deltacoronavirus sequences based on nucleotide alignment of ORF1ab+RdRp using the GTR+F+G4 evolutionary model chosen according to BIC. The outgroups represent sequences from *Alpha* (OQ540913.1), *Beta* (LC706864.1), and *Gamma* (MN509588.1) coronaviruses. The dark blue points show the sequenced genomes. B) Zoom of clade I showing the positioning of the newly sequenced deltacoronaviruses highlighted in bold tip labels. Tip point colors represent the host and support values are aLRT/ultrafast bootstrap.

## Discussion

Although *S. leucogaster* and *F. magnificens* breed in abundance along the coast of Brazil (Mancini, et al. 2016), no study has ever been conducted to investigate viruses in these species. Here, we present the results of 26 months of research on these two species within the MONA Cagarras, off the coast of Rio de Janeiro State. A total of 153 birds were tested for influenza A viruses, all of which were negative. However, 14.3% were found to be infected with a novel deltacoronavirus. To date, this is the first detection of coronaviruses in *S. leucogaster* and *F. magnificens*.

The genus *Deltacoronavirus* is classified into three subgenera: *Andecovirus*, *Buldecovirus*, and *Herdecovirus*, which include seven ratified species: Wigeon coronavirus HKU20 (*Andecovirus* subgenus), Bulbul coronavirus HKU11, Common Moorhen coronavirus HKU21, Coronavirus HKU15, Munia coronavirus HKU13, White-eye coronavirus HKU16 (*Buldecovirus* subgenus), and Night Heron coronavirus HKU19 (*Herdecovirus* subgenus) (Woo, et al. 2023). Six of these species represent groups of viruses found only in birds, while one species, Coronavirus HKU15 (*Buldecovirus*), represents a group of viruses detected in both birds and mammals, including pigs (Woo, et al. 2012). Globally, coronaviruses are distributed among wild birds, especially in orders such as Anseriformes, Charadriiformes, Columbiformes, Pelecaniformes, Galliformes, Passeriformes, Psittaciformes, Accipitriformes, Ciconiiformes, and Gruiformes, among others (Chu, et al. 2011; Wille and Holmes 2020; Chu, et al. 2022; Marchenko, et al. 2022). Although avian coronaviruses are generally associated with mild or even asymptomatic infections, some strains can cause more severe symptoms in birds, potentially leading to death (Cavanagh 2005).

Our findings reveal that *F. magnificens* and *S. leucogaster* harbor deltacoronaviruses. The virus appears to be the same in both species, suggesting the possibility of viral exchange, potentially through environmental contamination or direct transmission, as these two species coexist and nest in the same habitat (**Suppl. Figure 1)**. The longitudinal design of this study, spanning over two years, provides a unique opportunity to reflect on the potential dynamics of viral transmission within seabird colonies. The identification of deltacoronaviruses across different sampling periods and age groups (**Figure 2**) raises questions about possible sustained transmission or environmental persistence of these viruses within the colony. It indicates either sporadic introductions from external sources (e.g., migratory species) or intra-species maintenance, potentially facilitated by high-density nesting, parental care, or reuse of nesting sites (**Suppl. Figure 1**). Moreover, both *S. leucogaster* and *F. magnificens* are long-lived species, with life expectancies of at least 30 years and possibly up to 50 years (AnAge 2025). This extended lifespan increases the chances of lifetime exposure to diverse pathogens and may support the long-term maintenance of viral lineages within individuals or populations, even in the absence of frequent reintroductions. Such characteristics position these seabirds as potential long-term reservoirs or sentinels for viral surveillance in tropical marine ecosystems.

Very similar virus sequences have previously been identified in environmental samples (feces of *S. leucogaster*) from the São Pedro and São Paulo Archipelago, although only the partial RdRp was recovered (Gomes, et al. 2025). The genetic similarity between the viruses from these two locations, despite being over 1,000 km apart, suggests the possibility of pathogen exchange between these seabird populations, facilitated by migration or long-distance dispersal. While *F. magnificens* nests on islands throughout the Caribbean and in tropical areas along both coasts of Central and South America (Diamond and Schreiber 2002) (**Suppl. Figure 8**), some individuals may remain within their breeding range due to the lengthy breeding season, which can last for over one year. However, others, particularly pre-breeders and non-breeding birds, may disperse considerable distances from their breeding colonies (Diamond and Schreiber 2002). The GPS data from monitored frigatebirds have shown post breeding movements of up to 1,400 km (Weimerskirch, et al. 2006) with some individuals recorded dispersing as far south as the Arvoredo Island, off the coast of Santa Catarina (690 km away in a straight line), and to the northern part of Brazil (L.C., unpublished data). Similarly, *S. leucogaster* breeds in subtropical and tropical oceans between 25°S and 25°N (Nelson 1978), including the São Pedro and São Paulo Archipelago (Mancini, et al. 2016)(**Suppl. Figure 8**). While there is limited data on migration patterns, most birds in our study seem to be residents (Cunha L. personal observation). Nevertheless, data from other regions suggest that some individuals of *S. leucogaster* may disperse over distances of 800 to 2,900 km (Schreiber, et al. 2002), and even as far as 6,575 km (Kohno, et al. 2019). This evidence leads us to believe that the distribution of the novel deltacoronavirus is likely more widespread geographically and may also occur in other marine bird species.

Phylogenetic analysis indicates a close evolutionary relationship between the viruses we detected and those found in aquatic avian species from the orders Pelecaniformes, Anseriformes, and Suliformes (cormorants) in Asia and Australia, suggesting a potential common origin. However, the lack of comprehensive coronavirus sequence data from South America limits our ability to make more in-depth inferences. At this stage, it remains uncertain whether these viruses represent new species, as they form a distinct clade within the genus *Herdecovirus*. The detection of deltacoronavirus in 9.1% of *F. magnificens* and 18.4% of *S. leucogaster* suggests that these species are important hosts for the long-term maintenance of this virus. The prevalence of positive samples for deltacoronaviruses can vary significantly, ranging from approximately 0.3% to 50%, depending on temporal, seasonal, and spatial factors, as well as the detection methods used (Kim and Oem 2014; Wille, et al. 2016; Wille, et al. 2017; Milek and Blicharz-Domanska 2018; Wille and Holmes 2020). Even when the same detection method is employed, positivity rates can fluctuate between 0.95% and 15% (Chu, et al. 2011; Marchenko, et al. 2022). In addition to the detection of the virus in different individual samples in different years, when inspecting the genomic sequences recovered, we identified two independent in-frame deletion events in the Spike coding region that led to four amino acid deletions. These deletions in the N-terminal region of the Spike resemble Recurrent Deletion Regions (RDRs) extensively characterized in betacoronaviruses (McCarthy, et al. 2021), including in the SARS-CoV-2 variant of concern (Resende, et al. 2021). These in-frame adaptive deletions have been shown to emerge for several viruses under the selection pressure of neutralizing antibodies once they remove specific epitopes, rendering important neutralizing antibodies ineffective in binding to the viral Spike protein and hence neutralizing it (McCallum, et al. 2021). These findings suggest that either some *S. leucogaster* individuals sampled have been reinfected or developed long-term infection; in both cases, antibodies were likely elicited, and due to intrahost selection pressure, new lineages bearing in-frame deletions that likely affect neutralizing antibody binding emerged. Unfortunately, we were not able to resample the same individuals to test these alternative hypotheses. Moreover, additional analyses are warranted to further characterize Spike epitopes and the immunological response of these seabird species against the DeltaCoV found.

Several studies have revealed the significant role of wild aquatic and migratory birds, primarily from the orders Anseriformes and Charadriiformes, in the transmission of avian influenza viruses, highlighting the critical role of these bird species in virus dissemination (Webster, et al. 1992). The highly pathogenic avian influenza virus (HPAIV) subtype H5N1 of clade 2.3.4.4b has affected various species within the Sulidae family globally, including frigatebirds, shags, cormorants, gannets, and boobies (FAO 2024). The subtype H5N1 of clade 2.3.4.4b was first detected in Brazil on May 15, 2023, introduced by migratory birds from the south. Since then, it has affected 23 bird species and two marine mammal species. Among the birds, the most impacted species have been *Thalasseus acuflavidus* and *Thalasseus maximus* (MAPA 2023; Reischak, et al. 2023; de Araujo, et al. 2024). To date, there have been three confirmed cases of *S. leucogaster* dying from H5N1— one in Espírito Santo State and two in Santa Catarina State—and a single case of *F. magnificens* in Niterói, Rio de Janeiro State, all recorded in 2023 (MAPA 2023). Given these concerns, we closely monitored the colonies on the MONA Cagarras. Fortunately, no influenza virus was detected in any of the 153 seabirds sampled, nor were any antibodies found, indicating that the birds had not been exposed to any influenza virus so far. Additionally, no dead animals with suspected disease were found during our study. However, given their susceptibility to infection, continuous monitoring remains crucial.

Finally, additional viral taxa from the *Picornavirales* order were identified in four cloacal swab samples from *S. leucogaster* and *F. magnificnes.* The enterovirus phylogenetic tree revealed that the enterovirus sequences identified clustered within a clade with sequences from chimpanzees and birds. Interestingly, the identified here clustered together with other enteroviruses identified previously in feces of *S. leucogaster* from São Pedro and São Paulo Archipelago (PQ572693.1)(Gomes, et al. 2025). For sequences identified within the *Passerivirus* genus, we reconstructed one phylogenetic tree including the most similar sequences from the *Picornaviridae* family. The sequences from *S. leucogaster* clustered within a clade that was positioned together with other passeriviruses from thrushes. For *Caliciviridae* sequences, the CA50 partial RdRp was positioned as a basal branch of a clade containing other viruses such as Cryptolin_calicivirus_1 (WOC29232.1), Beihai_sesarmid_crab_virus_2 (YP_009333602.1), Wenzhou_picorna-like_virus_38 (APG78569.1), and Rudphi_virus_1 (AYN75566.1). These sequences were identified in other sea animals such as *Ruditapes philippinarum, Gastropoda,* and *Sesarmidae,* which are likely food sources of these seabirds.

## CONCLUSIONS

This study shows that marine birds from the MONA Cagarras are important hosts for deltacoronaviruses maintenance and transmission among seabirds, but not for influenza A viruses. Moreover, many other RNA viruses, such as enteroviruses, *Passerivirus*-like, and caliciviruses, were also characterized and warrant further studies to evaluate their impact on seabirds and sympatric species. Understanding the prevalence of pathogens in wild birds is important for assessing their role in the overall impact of these pathogens, including potential spillover events to domestic animals and humans. Continued and enhanced surveillance efforts are essential to monitor the presence of viruses, such as CoVs, IAVs, in avian populations and their potential threat to human and animal health. This concern becomes particularly relevant considering the ongoing spread of the highly pathogenic subtype of bird flu H5N1 across South America and the distribution overlap between seabirds and human populations.

## Ethical statements and permits

Fieldwork followed national guidelines and provisions of the ‘Sistema de Autorização e Informação em Biodiversidade’ (SISBIO, License numbers 73163-8, 73163-9). All animal procedures and veterinary assistance were in accordance with the ‘Conselho Nacional de Controle de Experimentação Animal’ (CONCEA, 2013), and approved by the ‘Comissão de Ética no Uso de Animais’ of Oswaldo Cruz Institute (License L-019/2021 and L-019/2021-A1) and registered in the ‘Sistema Nacional de Gestão do Patrimônio Genético e do Conhecimento Tradicional Associado’ (SisGen A3F2542).

## Acknowledgements.

We thank the Platform of Instituto Oswaldo Cruz, Fiocruz, Rio de Janeiro, Brazil, and the Genomic Platform—DNA Sequencing—RPT01A (Rede de Plataformas Tecnológicas Fiocruz). Special thanks to Sergio Jordão, the boat captain.

## Data availability statement

Raw reads generated in this study were submitted to the NCBI Sequence Read Archive (SRA) and are available under project number PRJNA1215064.

