## Supplementary figures and images for "Genomics insights reveal multi-year maintenance of a new *Deltacoronavirus* infecting Seabirds from the Cagarras Island Archipelago Natural Monument, Brazil"

### Suppl. Figure 1

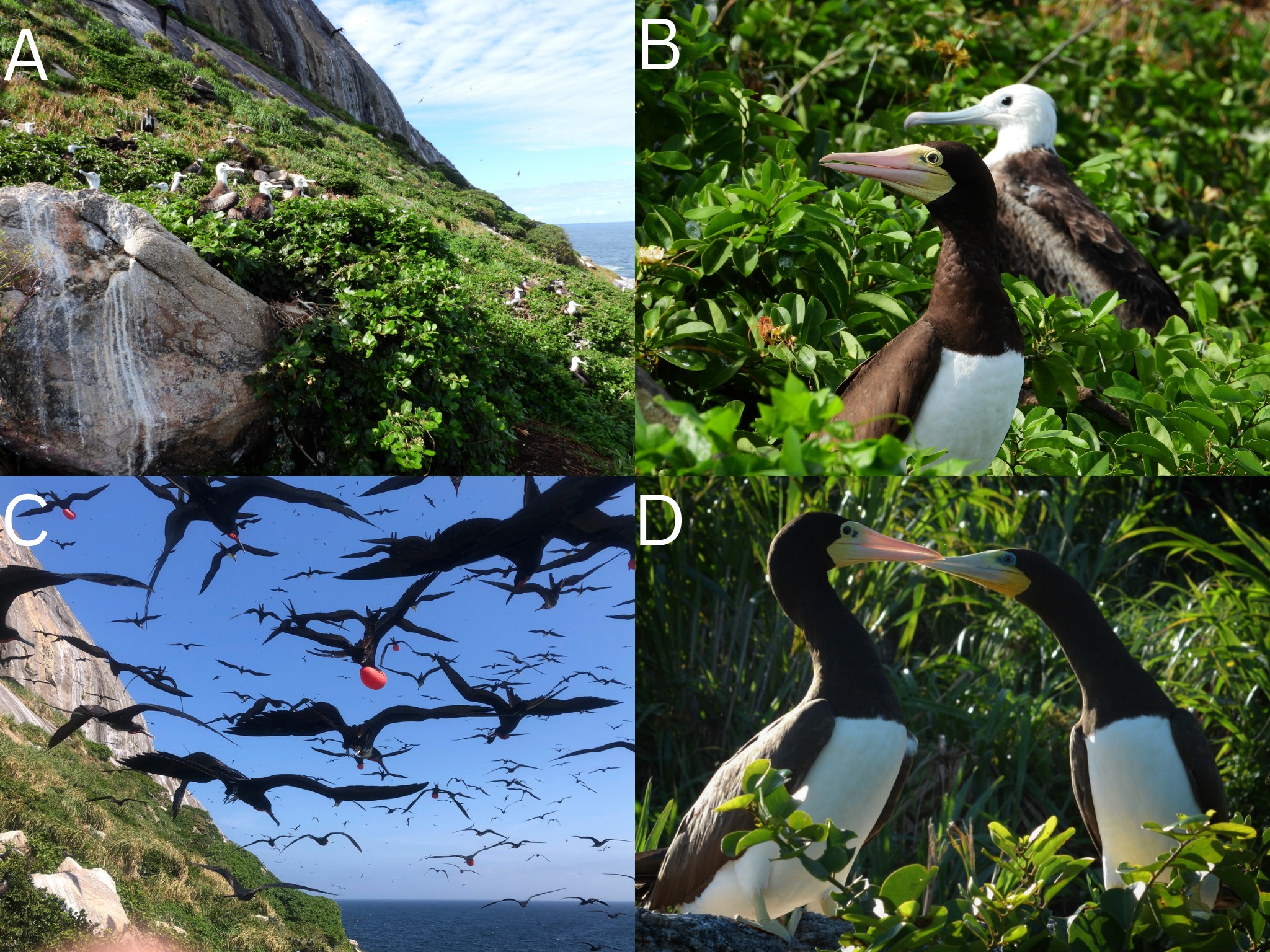

### Suppl. Figure 2

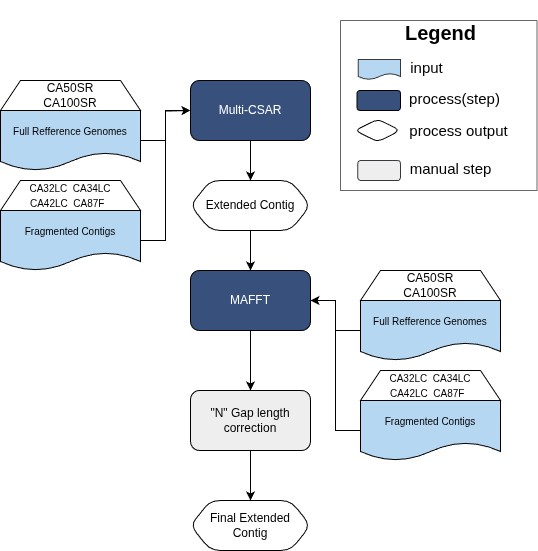

### Suppl. Figure 3

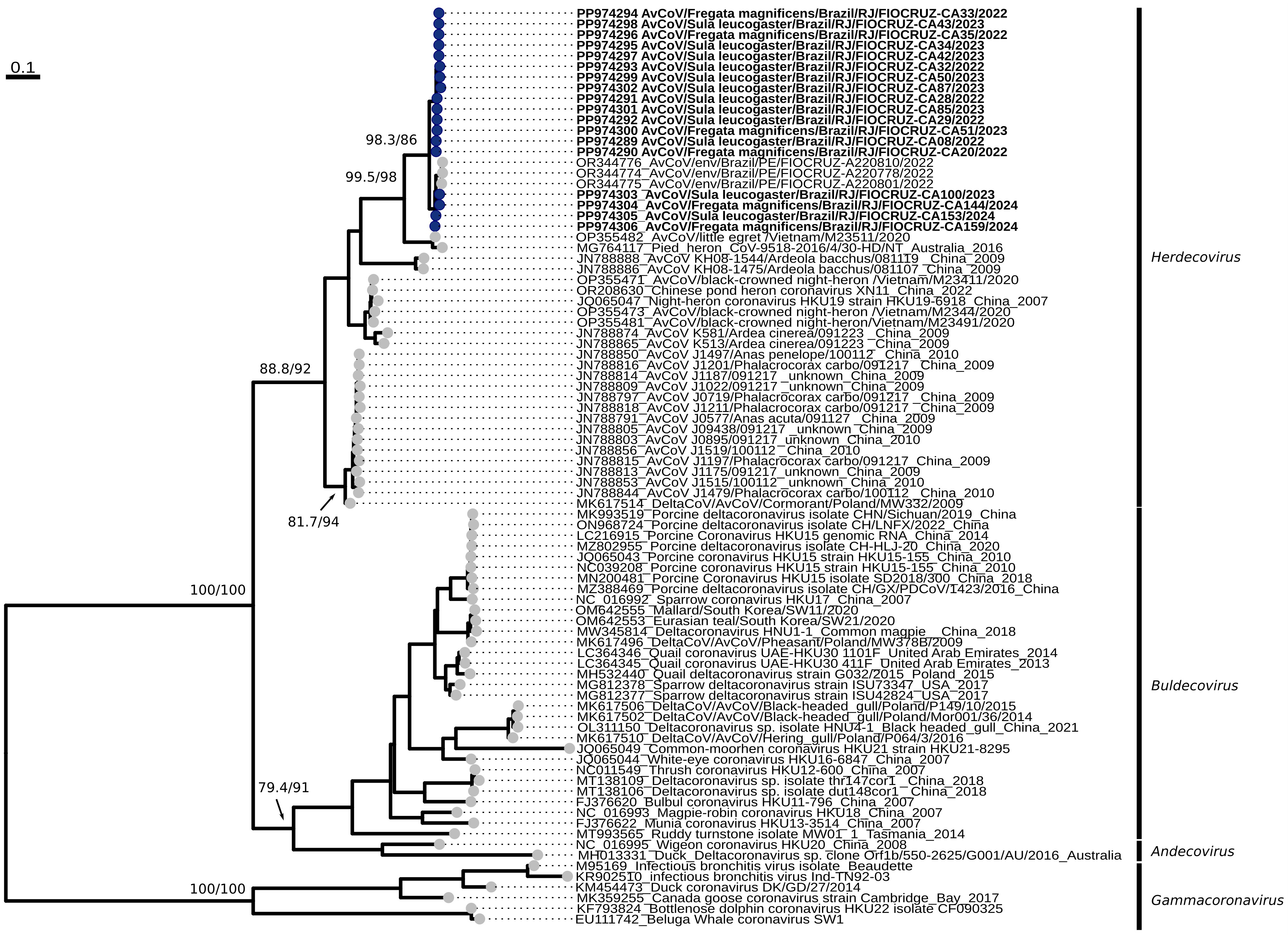

### Suppl. Figure 4

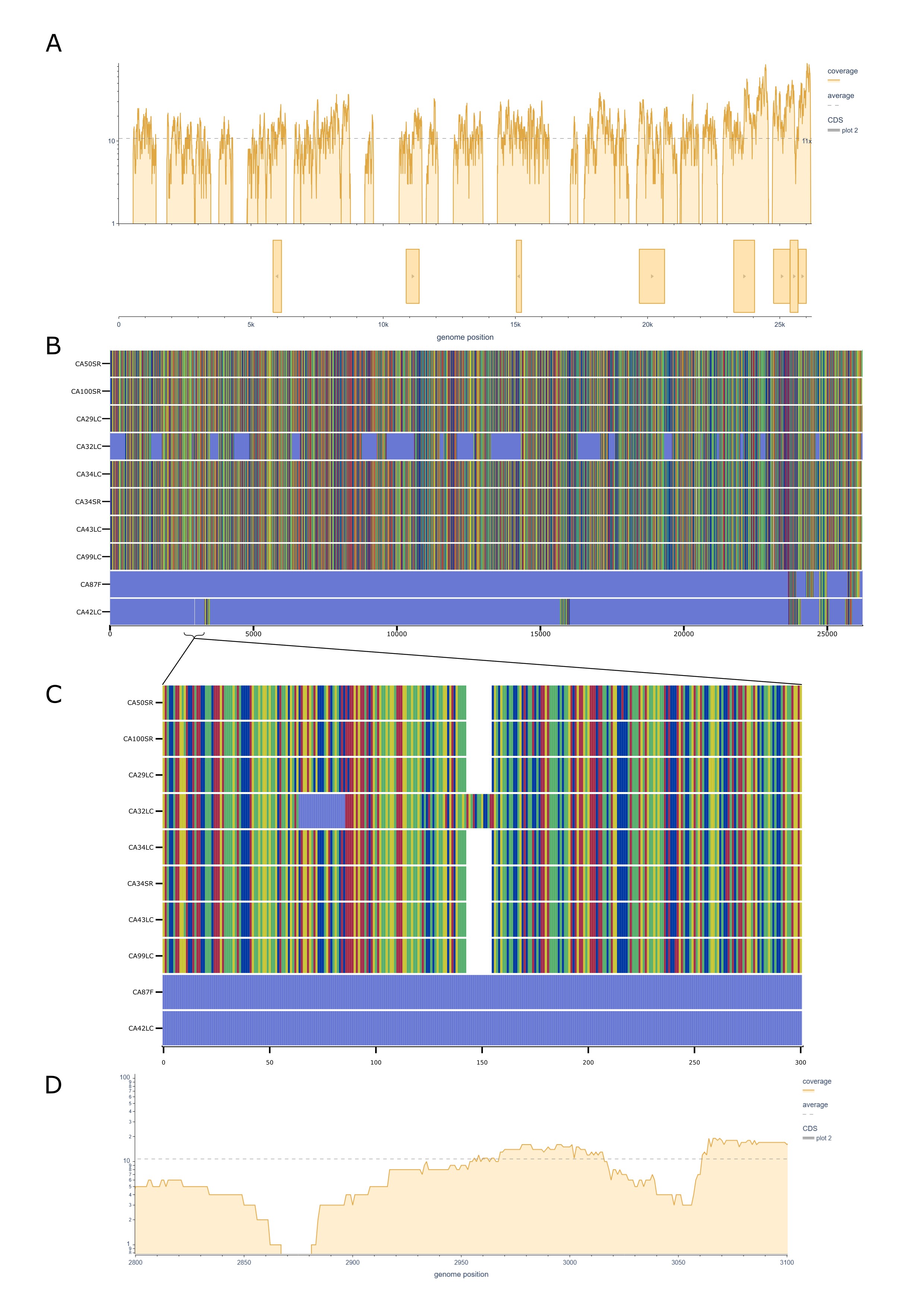

### Suppl. Figure 5

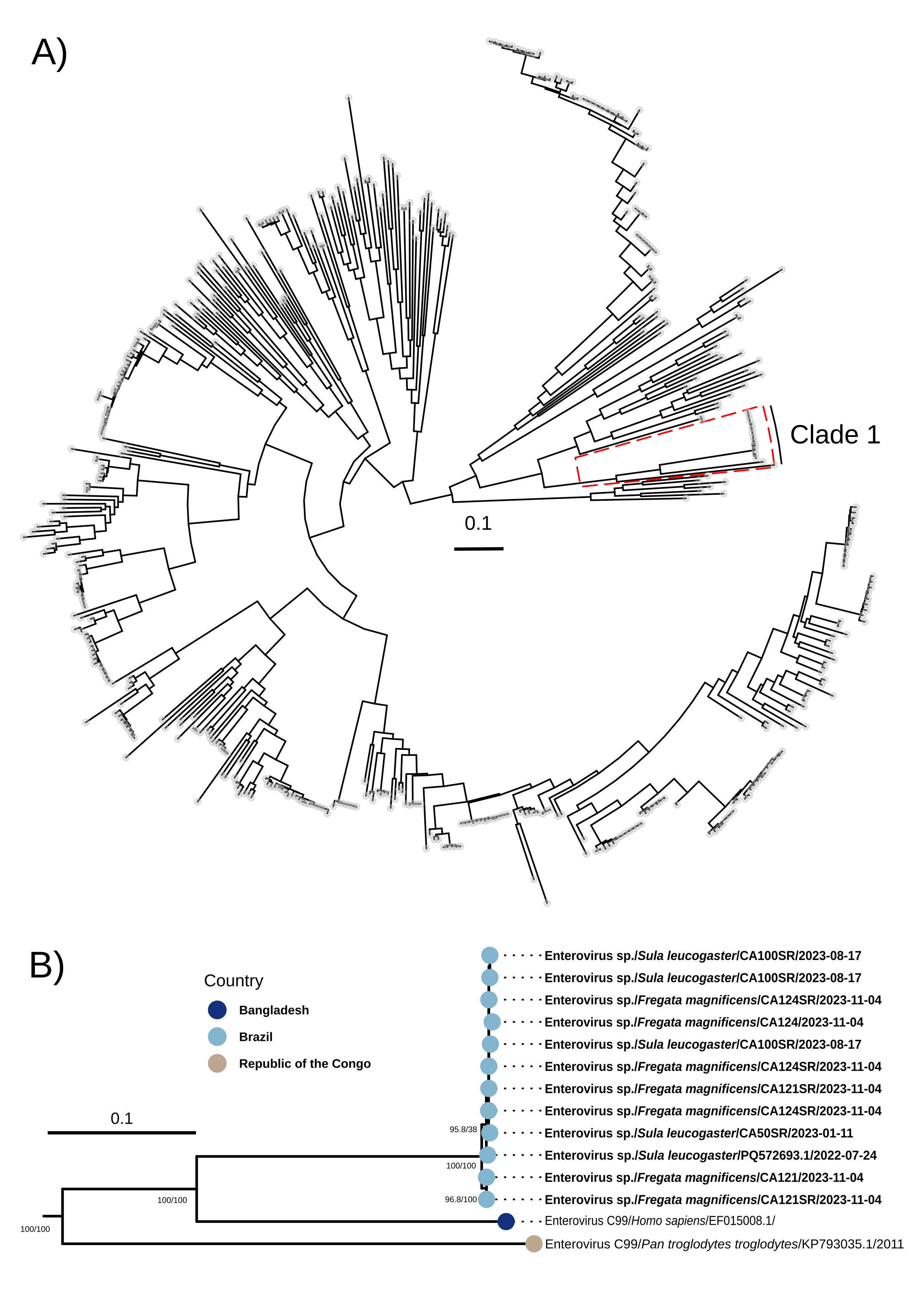

### Suppl. Figure 6

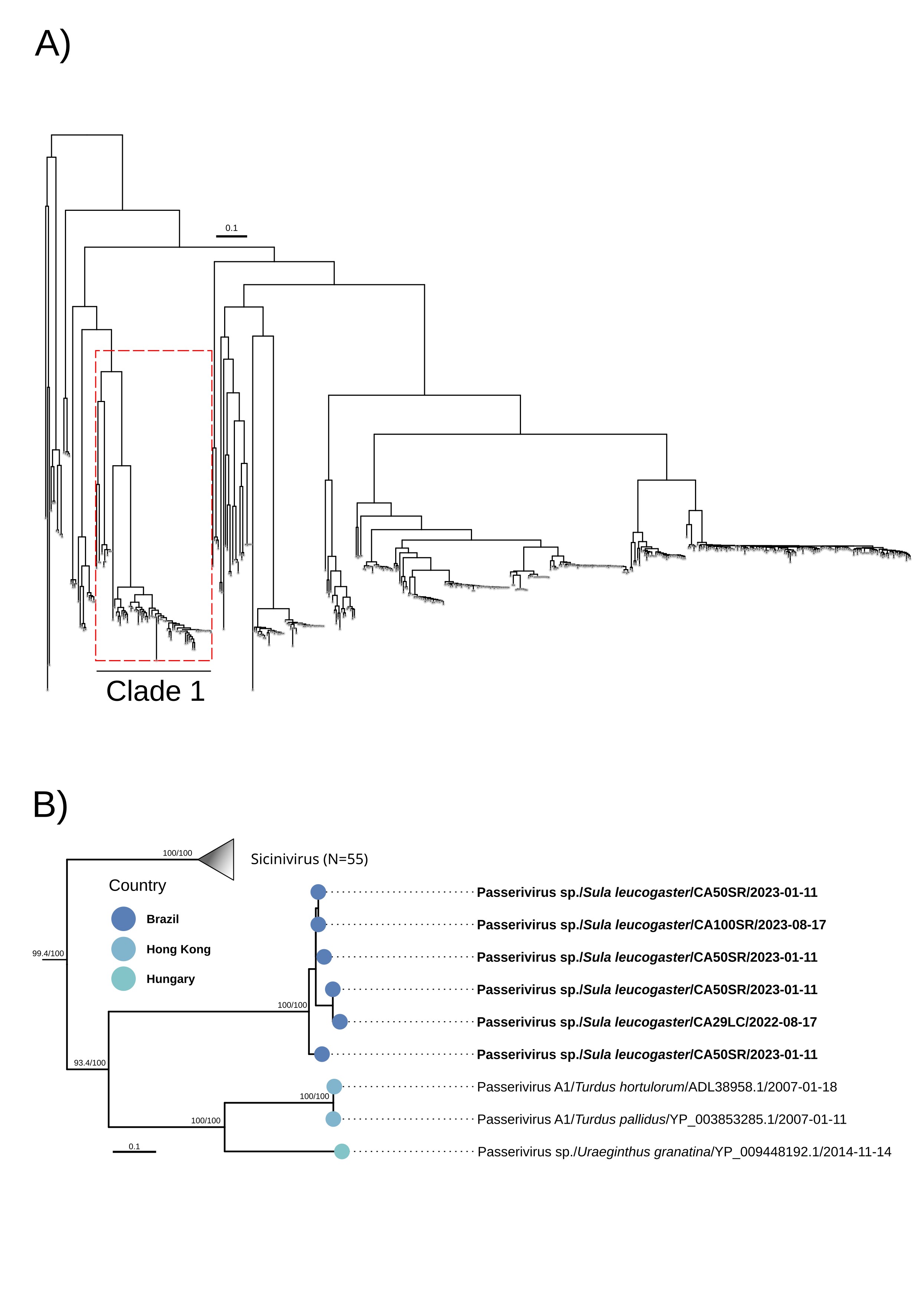

### Suppl. Figure 7

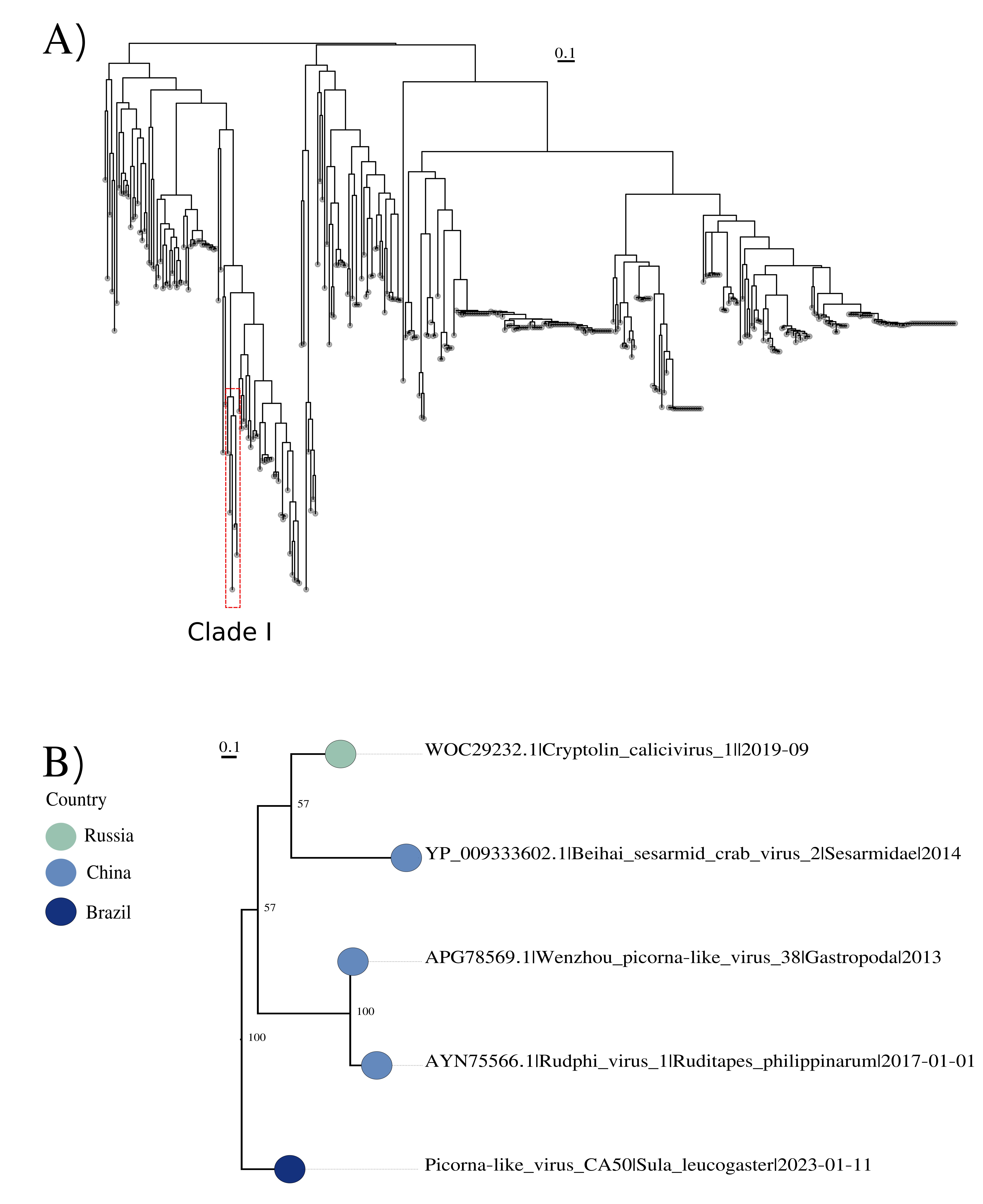

### Suppl. Figure 8

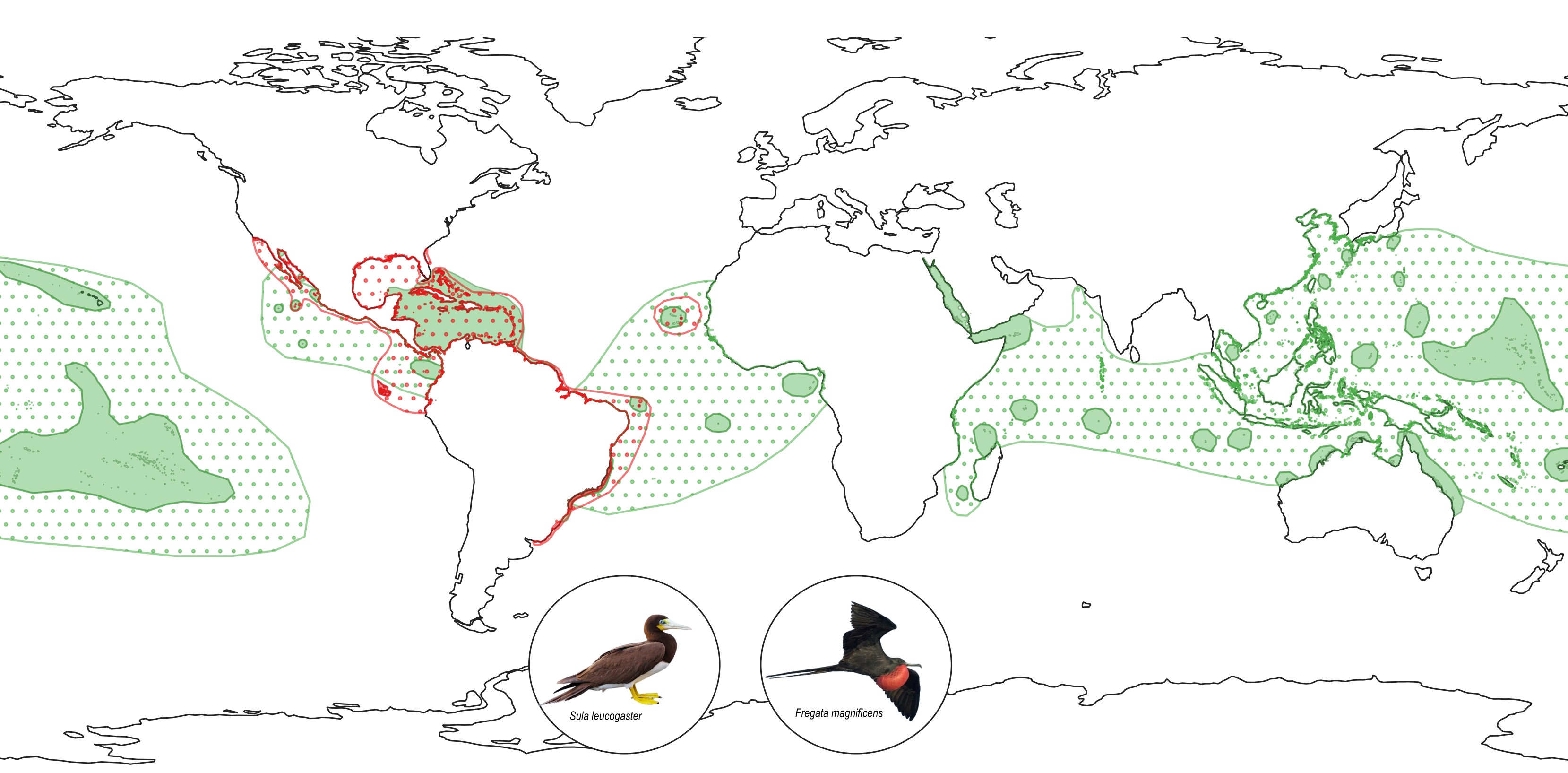
