## Supplementary material for "Genomics insights reveal multi-year maintenance of a new *Deltacoronavirus* infecting Seabirds from the Cagarras Island Archipelago Natural Monument, Brazil": Suppl. Table 2

**Suppl. Table 2**. Results from Chi-square tests conducted in R to assess the significance of variables - Species, Age, Sex, and Sample Type - on the presence of coronavirus. The analyses include comparisons across all variables and within species-specific groups to identify significant associations. Degrees of freedom (df).

| **Variables** | **Chi-squared** | **df** | **p-value** |
| --- | --- | --- | --- |
| **All variables** |  |  |  |
| Species vs Coronavirus | 0.051852 | 1 | 0.8199 |
| Age vs Coronavirus | 1.8667 | 2 | 0.3932 |
| Sex vs Coronavirus | 0.20741 | 2 | 0.9015 |
| Sample Type vs Coronavirus | 0.25846 | 1 | 0.6112 |
| *Sula leucogaster* |  |  |  |
| Age vs Coronavirus | 1.0057 | 2 | 0.6048 |
| Sex vs Coronavirus | 1.0057 | 2 | 0.6048 |
| Sample Type vs Coronavirus | 0.42328 | 1 | 0.5153 |
| *Fregata magnificens* |  |  |  |
| Age vs Coronavirus | 1.2857 | 2 | 0.5258 |
| Sex vs Coronavirus | 0.9 | 2 | 0.6376 |
| Sample Type vs Coronavirus | 0 | 1 | 1 |
